# Modified Bacterial Lipids Which Alter Membrane Surface Charge Reduce Binding of Antimicrobial Peptides

**DOI:** 10.1101/2020.04.24.057349

**Authors:** Patrick W. Simcock, Maike Bublitz, Flaviu Cipcigan, Maxim G. Ryadnov, Jason Crain, Phillip J. Stansfeld, Mark S.P. Sansom

## Abstract

Antimicrobial peptides (AMPs) initiate killing of bacteria by binding to and destabilizing their membranes. The multiple peptide resistance factor (MprF) provides a defence mechanism for bacteria against a broad range of AMPs. MprF reduces the negative charge of both Gram-positive and Gram--negative bacterial membranes through enzymatic conversion of the anionic lipid phosphatidyl glycerol (PG) to either zwitterionic alanyl-phosphatidyl glycerol (Ala-PG) or cationic lysyl-phosphatidyl glycerol (Lys-PG). The resulting change in membrane charge is suggested to reduce AMP-membrane binding and hinder downstream AMP activity. Using molecular dynamics to investigate the effects of these modified lipids on AMP-binding to model membranes, we show that AMPs have substantially reduced affinity for model membranes containing Ala-PG or Lys-PG. A total of ~7000 simulations are used to define the relationship between bilayer composition and binding for 5 different membrane active peptides. The reduction of degree of interaction of a peptide with the membrane is shown to correlate with the change in membrane surface charge density. Free energy profile (potential of mean force) calculations reveal that these lipid modifications alter the energy barrier to peptide helix penetration of the bilayer. These results will enable us to guide design of novel peptides which address the issue of resistance via MprF-mediated membrane modification.

## INTRODUCTION

An increasing number of pathogenic bacterial species are developing resistance to current antibiotics, paving the way to a post-antibiotic era in which previously treatable infections cause increased mortality^1-2^. Antimicrobial peptides (AMPs) are found in all kingdoms of life, in humans as part of the innate immune response^3-4^, and thus are a potential source for the development of novel antibiotics. Thousands of naturally occurring peptides have been identified^5^ and many synthetic ones have also been developed^6^. There is little sequence conservation between peptides found in different species^7^, thus providing a large data-set for the design of new peptides^8^.

AMPs have diverse sequences but are generally cationic and form an amphipathic α-helix, presenting distinct regions of hydrophobic and hydrophilic residues. These cationic peptides are rich in Arginine or Lysine^9^ and exhibit a strong correlation between charge (typically in the range of +2 to +9) and antimicrobial activity^10^. AMPs selectively target bacterial membranes^11^. Once adsorbed onto a membrane they can induce a range of effects^12^ including membrane deformation and disruption, membrane depolarization, and clustering of charged lipids, and various modes of pore formation^13^ alongside a detergent-like mechanism^14^ Many of these effects have been observed experimentally for the canonical AMP magainin (from *Xenopus laevis*)^15^. NMR studies have shown that at low concentrations magainin adsorbs onto lipid bilayer membranes, interacting with lipid head-groups^16-17^ and reducing local membrane thickness^18^. At high Peptide/Lipid (P/L) ratios (> 1/30), peptides move from a surface-bound to a trans-membrane orientation^19^ and form a toroidal pore in the bilayer^20^. At this P/L, dissipation of the transmembrane potential^21^ and membrane leakage^22^ of cell contents are observed. Mechanisms for AMPs depend not only on peptide structure and charge, but also on membrane lipid composition^23-24^.

Some bacteria have developed resistance to AMPs by changing their cell surface electrostatics^25^. Certain Gram-positive bacteria (e.g. *Staphylococcus aureus, Bacillus subtilis* and *Clostridium perfgrinens*^26^) can modify their lipids to change the membrane surface charge through the action of the multiple peptide resistance Factor (MprF) protein^27^. This mechanism also forms an additional defence mechanism in some Gram-negative bacteria (e.g. *Pseudomonas aeruginosa*^28-29^ and *Rhizobium tropici*^30^) and also in *Mycobacterium tuberculosis*^31^.

MprF is a transmembrane protein that acts as both an enzyme (a synthase) and as a membrane transporter (a flippase) for modified phosphatidyl glycerol (PG) lipid species. PG is a major anionic lipid in bacterial membranes. MprF consists of a C-terminal enzymatic domain responsible for aminoacylating the headgroup of PG and a hydrophobic N-terminal domain responsible for flipping the modified lipid to the outer leaflet of the cytoplasmic membrane^32^. MprF can transfer lysine obtained from Lysyl-tRNA to transform PG into a positively charged lipid Lys-PG^33^ (SI Fig. S1). Alanine is also a common substrate used by MprF paralogs to produce zwitterionic Ala-PG^34^. The presence of MprF correlates with a reduction in the activity of AMPs against resistant bacteria. Incorporation of Lys-PG into artificial membranes significantly impairs the formation of membrane defects caused by AMPs^35^. MprF is critical for production of these modified lipids^36-37^, which also provides protection against peptide antibiotics such as daptomycin^38-41^ (a lipo-peptide) and vancomycin^42-43^ (a glyco-peptide) in addition to AMPs^44-45^. This is of clinical importance, as these antibiotics are reserved for combatting pathogens already resistant to many other drugs, highlighting the importance of understanding the mechanism and role of MprF.

Molecular dynamics (MD) simulations have been widely used to explore possible mechanisms of action of AMPs^46-48^, and in particular their interactions with lipid bilayer membranes. MD simulations have shown individual peptide molecules will rapidly adsorb onto the membrane surface^49-51^, interacting with the lipid head-groups and inserting into the membrane core when multiple peptides are present, with a preference for negatively charged membranes^52^. Further simulations supported by experiments suggested that peptides insert into membrane bilayers forming transmembrane or monolayer pores depending on their orientation in the bilayer^53-54^. Peptides self-inserting into the membrane have been shown to form disordered toroidal pores^49 22^. MD simulations have also been used to explore various other possible peptide pore structures^55-57^. Coarse-grained (CG) MD simulations have also shown that peptides will adsorb onto membranes^58^ with a preference for negatively charged surfaces and that pores pre-formed in atomistic simulations remain stable when mapped to a CG representation^59^.

Here, we use CG MD to examine the effect of the two types of MprF-modified lipids, Ala-PG and Lys-PG, on peptide binding to bilayer membranes. We apply this methodology to five α-helical peptides of varying charge. A total of ~7000 simulations (see SI Tables S1 to S3) are used to define the relationship between bilayer lipid composition and AMP binding. Free energy profiles are calculated to show that MprF-mediated lipid modifications increase the energy barrier of single peptide penetration of the bilayer membrane.

## METHODS

Simulations were carried out using the molecular dynamics package GROMACS 2018.6^60-61^ using the coarse-grained Martini 3 force field^62-63^. Membrane models were generated using the INSANE^64^ methodology. After initial setup, all systems were energy minimised using the steepest descents approach.

All unbiased simulations underwent 5 ns of NVT and 10 ns of NPT equilibration with the peptide restrained in its initial position before launching a 1 μs unrestrained production run. All simulations were run with a timestep of 20 fs. Temperature was coupled at 310 K using a velocity rescale thermostat^65^ with a coupling time constant of 1 ps. Pressure coupling was carried out by a semiisotropic Parrinello-Rahman barostat^66^ at a reference pressure of 1 bar, compressibility at 3×10^−4^ bar^−1^ with a coupling time of 12 ps. Electrostatic interactions were calculated using a reaction field potential with a cut-off of 1.1 nm. Van der Waals interactions were calculated using a Verlet cut-off scheme for beads within 1.1 nm of each other and the potential switched from 0.9 nm.

Free energy landscapes were determined by calculation of potentials of mean force (PMFs) with a collective variable (i.e. reaction coordinate) generated by pulling a peptide across the membrane. Bilayers were generated with ~170 lipids (of a single species – see below and SI Table S3) in each leaflet in a 10 x 10 x 25 nm^3^ box. The peptide was pulled through the membrane with the N terminus first, with the *z* coordinate of the peptide ranging from −5 nm to +5 nm from the bilayer centre. Peptides were pulled through the membrane N terminus first, as this has been suggested to be a preferred binding pose^67-68^. Starting conformations were generated from the pull every 0.1 nm and simulated for 1 μs.^67-68^A restraining force of 1000 kJ mol^−1^ nm^−2^ was employed in the *z* direction during each simulation window. Free energy landscapes were built using the weighted histogram analysis method^69^.

Analysis was carried out using GROMACS tools and MDAnalysis^70^. VMD^71^ and PyMOL^72^ were used for molecular visualisation and rendering.

## RESULTS & DISCUSSION

### Peptides and Lipid Bilayers Studied

The overarching aim of this study is to test whether AMP resistance via MrpF modification of bacterial membranes can be explained by a change in the initial interaction of an AMP molecule with the membrane surface. To this end we selected five α-helical peptides which are all amphipathic but which differ in their net charge and their antimicrobial activity (Fig. 1): magainin 2 (Mgn2) is a canonical cationic AMP^15, 73^; cp5 and cp7 are synthetic cationic AMPs with relatively simple sequences^74^}; and amhelin (Amh) is a *de novo* peptide designed to be an archetypal model of AMPs^75^. A negatively charged amphipathic peptide which lacks antimicrobial properties (nAMP) is included as a control. Lipid bilayer compositions were chosen to provide simple models of both unmodified bacterial membranes (PE:PG ~2:1) and models of MprF-modified membranes containing either Ala-PG or Lys-PG in place of PG. The compositions of these membranes were varied to model bacteria which different levels of MprF-mediated lipid modification (see SI Tables S1 and S2).

**Figure 1:**
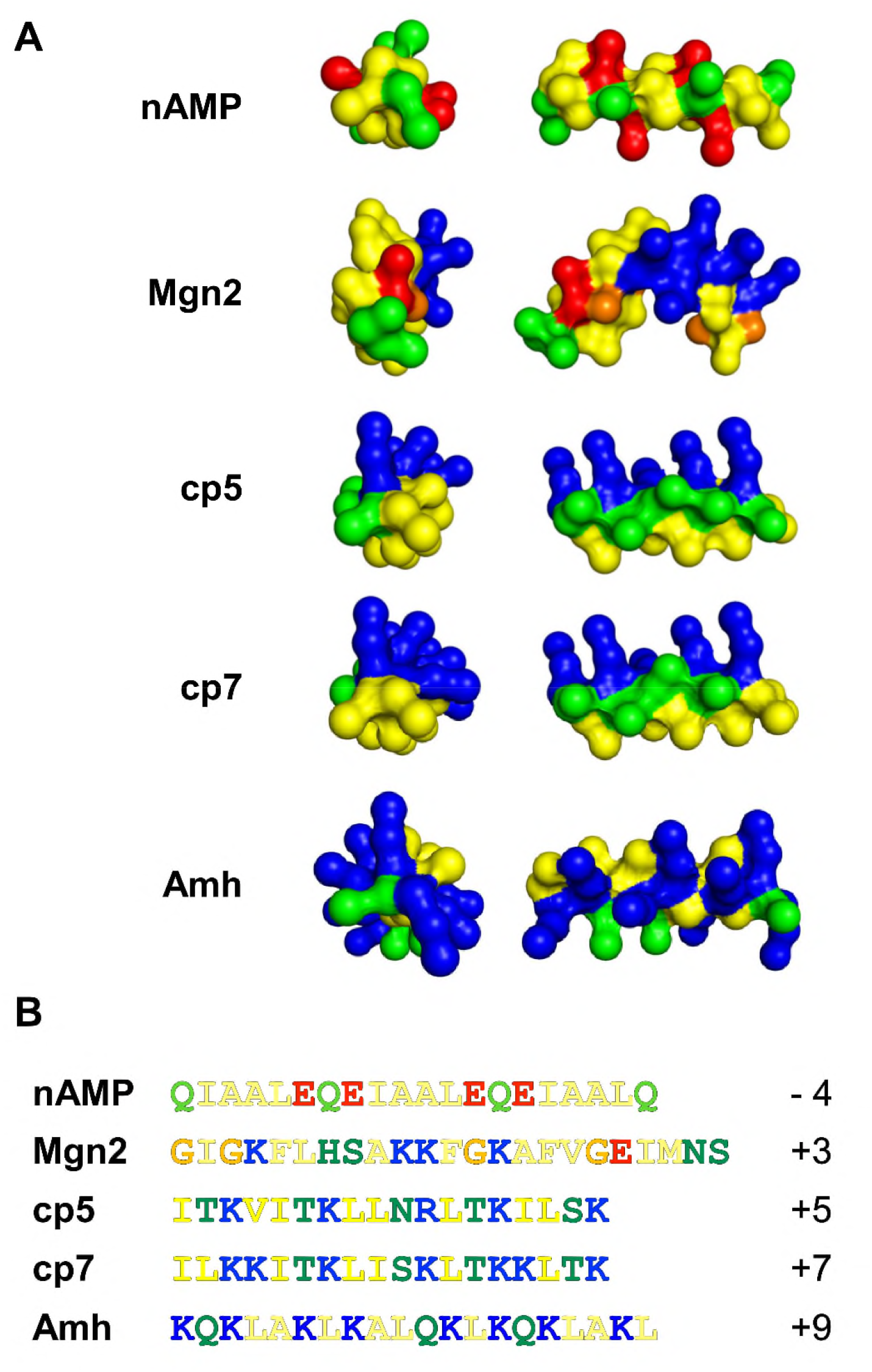
**A** Structures of peptides shown in coarse-grained representation with cationic amino acid residues in blue, anionic residues in red, polar uncharged residues in green, and hydrophobic residues in yellow. All peptides were modelled as continuous α-helices, apart from magainin 2 (Mgn2) for which the NMR structure determined in the presence of DPC micelles (PDB ID 2MAG) was employed. **B** The sequences and net charges of the peptides.

### Simulations reveal reduction of peptide binding due to lipid modifications

We first investigated whether membranes containing modified PG lipids showed reduced binding by AMPs compared to membranes in which only PG and PE lipids were present. An asymmetric lipid bilayer system was constructed. One leaflet of the bilayer contained PG, mimicking an unmodified (i.e. anionic) bacterial membrane (Fig. 2A). The other leaflet of the bilayer contained either Ala-PG or Lys-PG, representing a modified membrane. It is important to note that because of the use of periodic boundaries in the simulation, a peptide molecule initially placed in the aqueous phase about ~15 nm from the bilayer mid-plane (Fig. 2B), was able to freely access either face of the asymmetric membrane (Fig. 2C). Fifty simulations, each of a duration of 1 μs, were run for each peptide-membrane system, and the position of the peptide relative to the two bilayer surfaces vs. time was tracked (Fig. 2D,E). Peptide binding levels to each leaflet were calculated as the percentage of simulations that resulted in the peptide interacting with a given leaflet. Binding was defined to have taken place when all peptide backbone beads were within 0.6 nm of the closest phospholipid phosphate.

**Figure 2:**
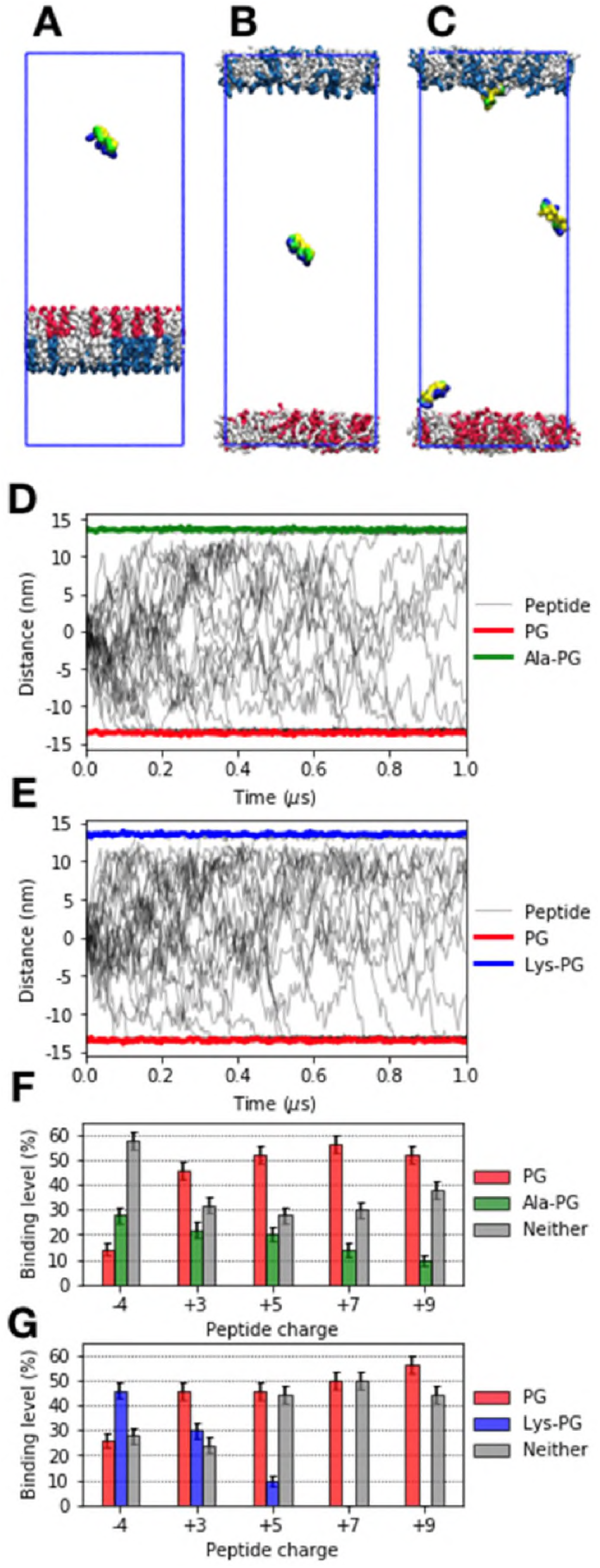
**A** Peptides were placed in a simulation box along with a bilayer membrane with an asymmetric lipid composition. The bilayer contained ~225 lipids per leaflet in a 12 x 12 x 30 nm^3^ box. **B** The membrane is split across periodic boundary conditions. A peptide molecule was placed in the centre, ~13 nm from each leaflet surface. **C** Over 50 repeats, the peptide *z*-position is recorded and the binding to each leaflet is determined. Trajectories of the +5 peptide with the **D** Ala-PG system and **E** Lys-PG systems are shown. **F** Binding levels for peptides with either the PG:PE, Ala-PG:PE leaflets or neither. **G** Binding levels for peptides with either the PG:PE, Lys-PG:PE leaflets or neither. Binding levels are defined as the percentage of simulations that result with the peptide binding to that specific leaflet. Errors are calculated through bootstrapping analysis.

Comparing the 5 peptides (see Fig. 1) and the two lipid modifications (Ala-PG, Fig. 2F and Lys-PG, Fig. 2G) our results demonstrate that the anionic PG leaflet is greatly preferred by all four cationic peptides relative to the zwitterionic Ala-PG and cationic Lys-PG leaflets. Furthermore, the more highly positively charged peptides show a greater preference for PG over the modified membranes, such that for peptides cp7 (charge +7) and Amh (charge +9) there were no detectable binding events with the Lys-PG leaflet. Comparison between the Ala-PG and Lys-PG systems showed that the Lys-PG modification had a more drastic effect than the Ala-PG modification. In contrast, the anionic control peptide nAMP displayed enhanced interaction with the Ala-PG and Lys-PG leaflets.

### Peptide binding is modulated by the lipid bilayer composition

We have seen that complete replacement of PG in a model bacterial membrane with either of the modified variants, Ala-PG or Lys-PG, leads to reduced selective binding of cationic peptides. However, *in vivo* membranes of bacteria expressing MprF are expected to contain both PG and Ala-PG and/or Lys-PG. The fractional composition of the modified lipids is believed to be dependent on bacterial strain and growth conditions, such that an acidic environment results in a larger number of modified lipids^29-30^. We therefore investigated peptide binding to bilayers composed of mixtures of PE and both PG and modified-PG lipids, thus defining how the bilayer composition of modified lipids affected the ability of AMPs to bind to the membrane.

Symmetric membranes of three components (PE:PG:Lys-PG or PE:PG:Ala-PG) were generated with lipid compositions ranging between PE (100%), PG:PE (50%:50%) to Lys-PG (or Ala-PG):PE (50%:50%) as defined on a triangular grid (Fig. 3A) with 10% increments in lipid composition between adjacent grid points (see also SI Table S2). This approach defined a lipid composition space for which AMP binding was investigated via 25 x 1 μs simulations for each grid point (i.e. each lipid composition; Fig. 3B). The binding level for all 5 peptides was thus determined by the total number of interaction events with the bilayer, for both the PE:PG:Lys-PG and the PE:PG:Ala-PG systems (Fig. 4A).

**Figure 3:**
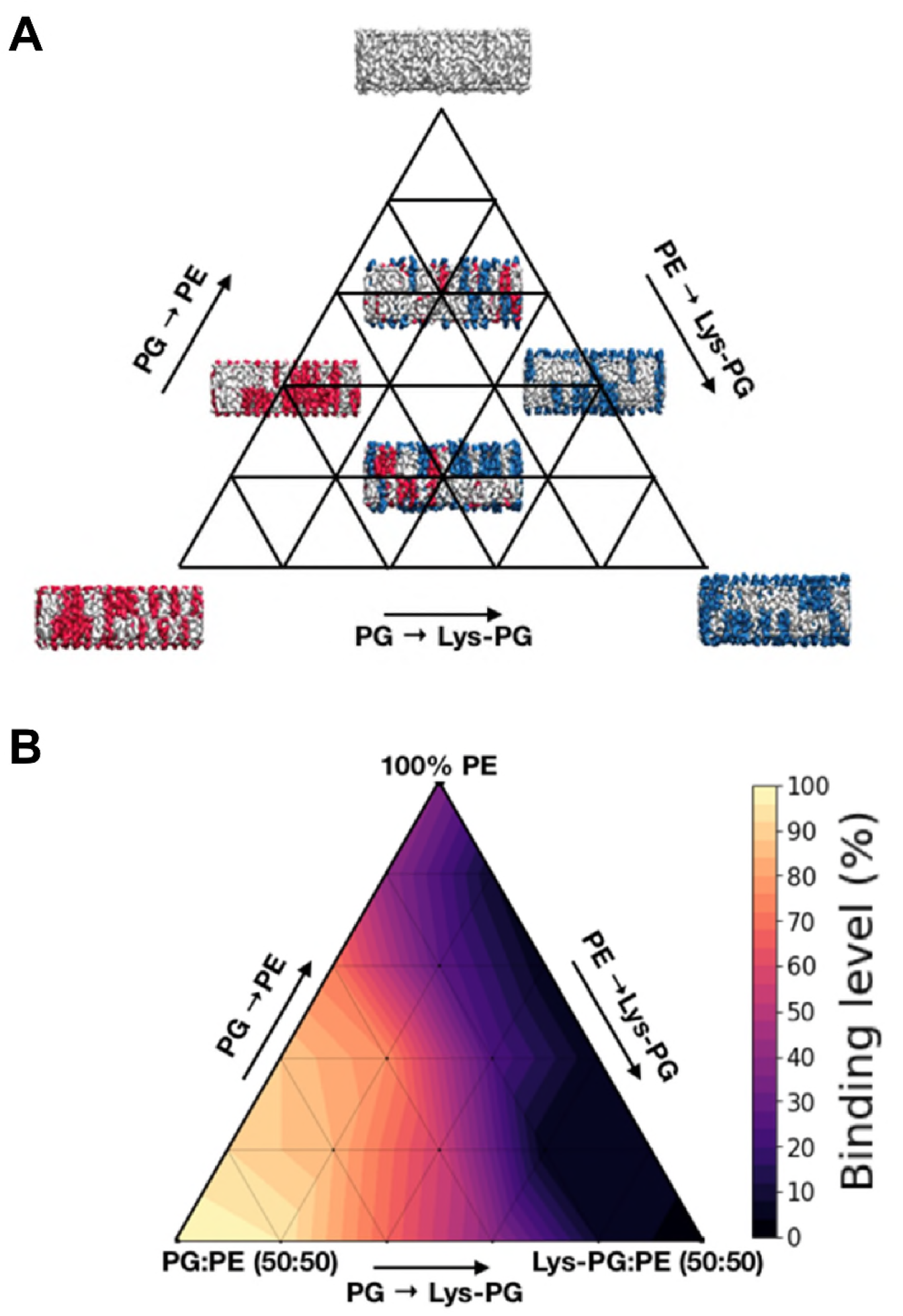
**A** Construction of lipid composition space to be investigated. Lipid compositions vary between 100 % PE, 50 % PG 50% PE and 50% Lys-PG or Ala-PG:PE. Membrane composition is varied by 10% at a time. Also see SI Table S2. **B** Peptide binding level vs. lipid composition space for a PG/PE/Lys-PG system with the Amh peptide. Binding level is calculated as the percentage of simulations in which the peptide binds to either leaflet of the membrane. A binding level of 0% (black) corresponds to no binding in any of the 25 repeats for that lipid composition, and a level of 100% (bright orange) represents every simulation resulting in binding.

All of the cationic peptides exhibited greater levels of binding when PG was present in the membrane, whilst replacement of PG by Lys-PG resulted in a reduction in binding. For the most positively charged peptides, i.e. cp7 and Amh, binding decreased from ~100% to 0% when comparing membranes containing 50% PG to those containing 50% Lys-PG. The replacement of PG with Ala-PG also resulted in decreased cationic peptide binding, though this effect is less than that of Lys-PG, especially for the most positively charged peptides. In contrast, nAMP binding increased from ~30% with a 50% PG membrane to ~ 100% binding with the 50% Lys-PG membrane.

Broadly speaking, for any given peptide the binding level correlates with the membrane charge density, with the sign and strength of the correlation depending on the charge on the peptide (Fig. 4BC). Thus, binding levels for nAMP show a positive correlation with increased (i.e. less negative, more positive) membrane surface charge density caused by lipid modification for both Ala-PG (Fig 4B) and Lys-PG (Fig 4C). The three most positively charged peptides (cp5, cp7 and Amh) all show a strong negative correlation with increasing (positive) membrane charge density, whilst Mgn2 shows intermediate behaviour. Thus, whilst Mgn2 exhibits a decrease in binding upon the inclusion of Ala-PG and Lys-PG, its binding level rarely falls below 50% regardless of the lipid composition, suggesting less highly charged peptides may be more resilient to changes in the membrane electrostatic environment.

**Figure 4:**
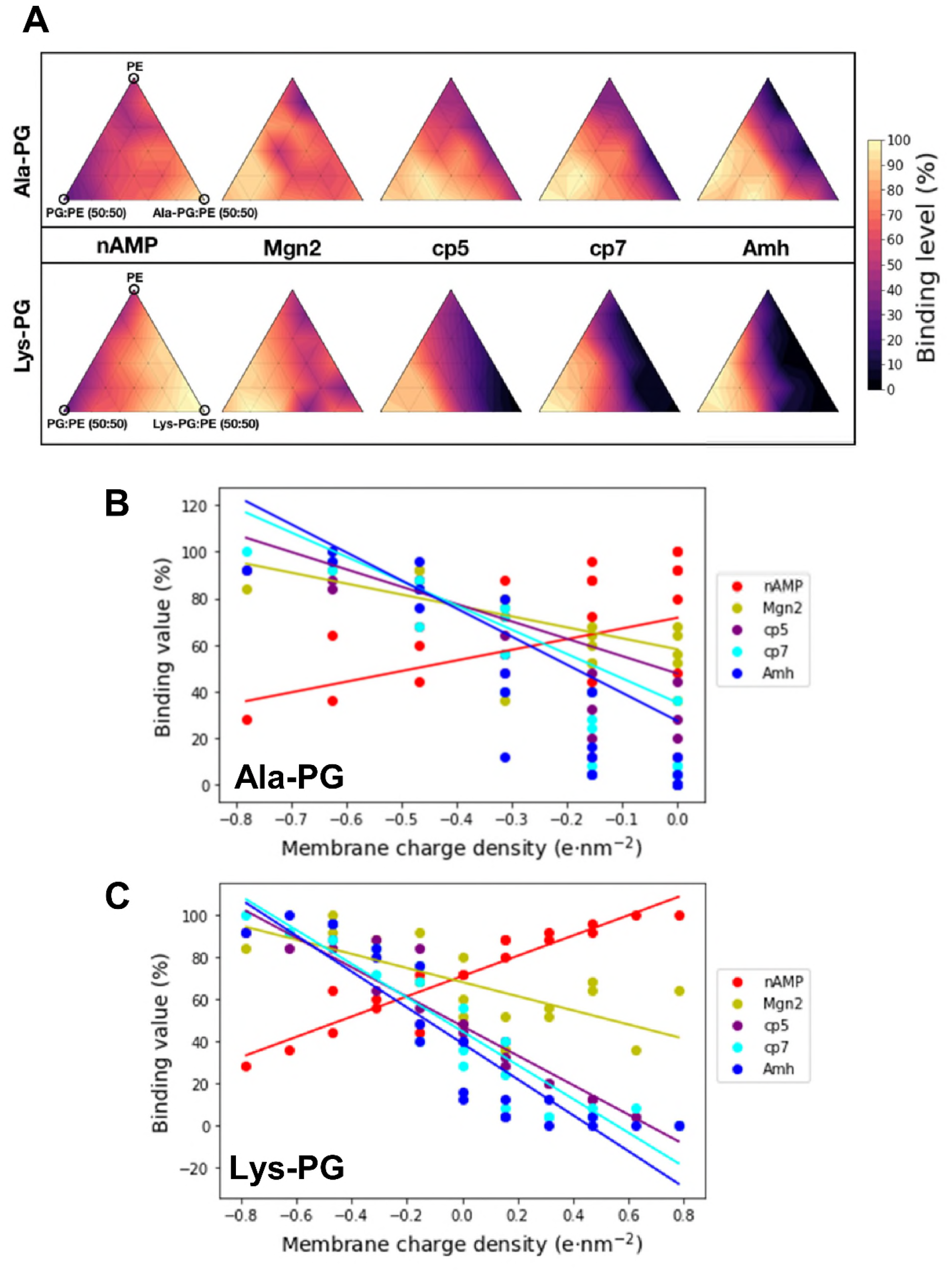
**A** Peptide binding level maps for peptides in membranes of varying lipid compositions. The binding in lipid composition is for each peptide is determined from 900 1 μs simulations. This data together represents 4.5 ms of data. **B,C** Correlation between the level of binding and the charge density of the bilayer for B Ala-PG containing bilayers and **C** Lys-PG containing bilayers, corresponding to the binding level maps shown in **A**.

### Free energy landscapes for single peptide molecule interactions with lipid bilayers

Free energy landscapes for the interaction of single peptide molecules with bilayers of differing lipid compositions can provide a mechanistic understanding of the trends analysed above. To this end free energy landscapes were calculated using Potential of Mean Force (PMF) calculations with a reaction coordinate corresponding to the distance between the centre of mass of a single peptide molecule and the centre of mass of the lipid bilayer. For every position along this reaction coordinate the peptide was allowed to rotate relative to the bilayer normal so as to fully sample all possible orientations relative to the membrane.

PMFs were generated for the interaction of the five different peptides with lipid bilayers containing a single species of lipid (i.e. PE, PG, Lys-PG or Ala-PG; see SI Table S3). Thus, a total of 20 PMFs were estimated, as shown in Fig. 5. In these PMFs the initial position of the peptide was with its N terminus oriented towards the bilayer. The peptides were otherwise free to rotate. Analysis of the tilt angle between the peptide helix axis and the bilayer normal (SI Fig. S2) shows that the peptides rotate freely whilst in the aqueous region outside the bilayer, adopt a tilt angle of ~90° (corresponding to parallel to the bilayer surface) when at the water/bilayer interface, before switching to the ~180° whilst the helix spans the bilayer (centred around a reaction coordinate value of ~0 nm). The two halves of the PMF are approximately equivalent, with the small difference between the two corresponding to whether the N terminus or the C terminus of the peptide is approaching and/or penetrating the bilayer surface (SI Fig. S3).

**Figure 5:**
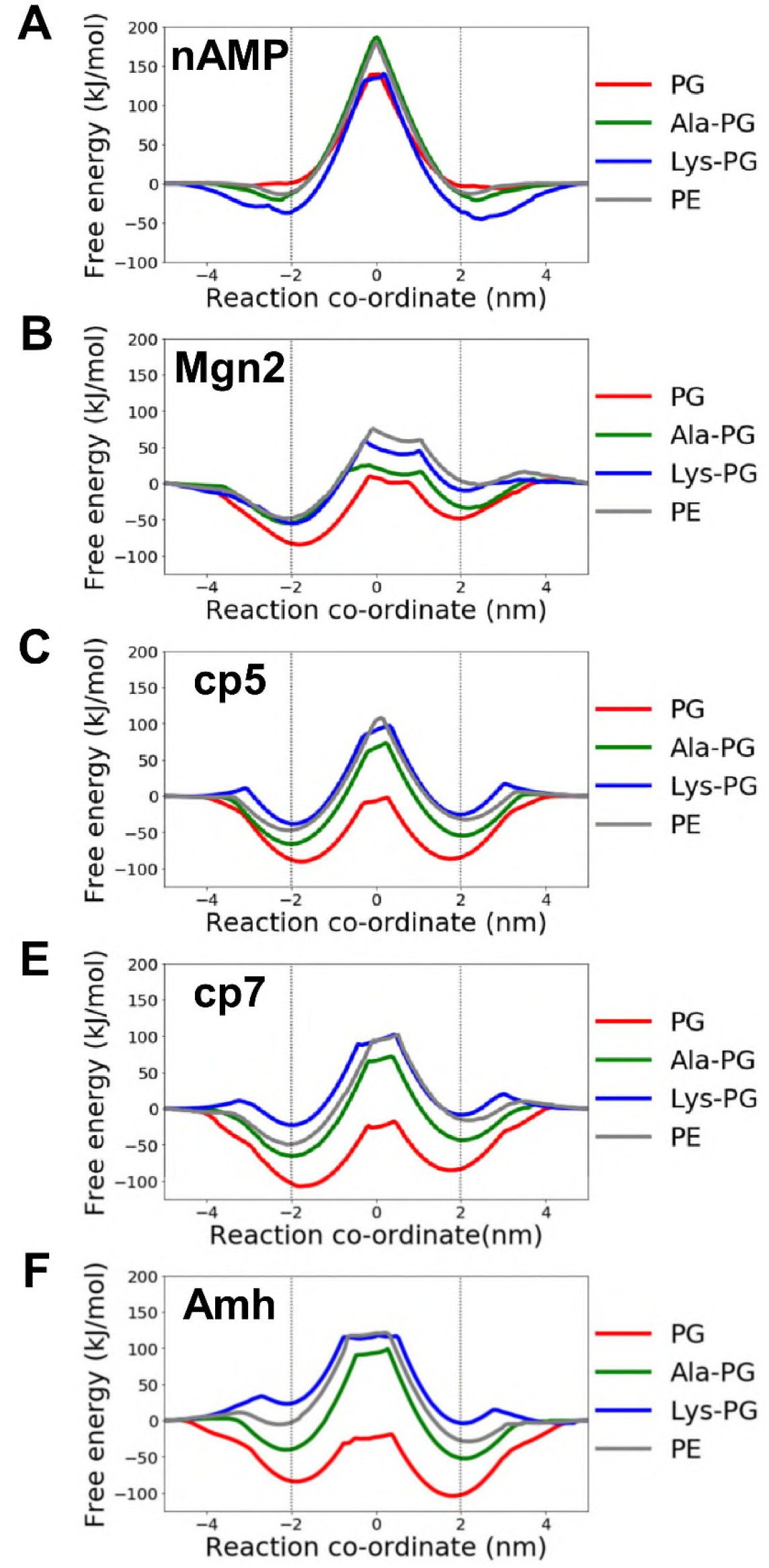
Potentials of mean force (PMFs) for single peptide helices with bilayers composed on PG (red), Ala-PG (green) Lys-PG (blue) or PE (grey). See text for details.

The general shape of the PMFs is as anticipated (Fig. 5), with a central barrier corresponding to a transmembrane orientation of the peptide helix, with free energy wells either side of this corresponding to the peptide lying at the bilayer/water interface, interacting extensively with the lipid headgroups. The depth of the free energy well for each peptide, corresponding to the free energy difference for transfer of the peptide from the aqueous phase to the peptide bound at the water/membrane interface, can be seen to depend on both the lipid species and the charge on the peptide Fig. 6A). From these data we may calculate *ΔΔG_WELL_*(PG-KPG *or* PG–APG) which corresponds to the change in well depth when a PG membrane is changed to either Lys-PG or Ala-PG. Plotting *ΔΔG_WELL_* as a function of peptide charge reveals a linear relationship for Lys-PG and for Ala-PG, in both cases passing though the (0,0) origin (Fig. 6B). This demonstrates the importance of peptide charge and the stronger effect of PG being replaced by Lys-PG than by Ala-PG. Note that these graphs show the effect on the free energy of peptide binding of complete replacement of a PG membrane by either Lys-PG or Ala-PG.

**Figure 6:**
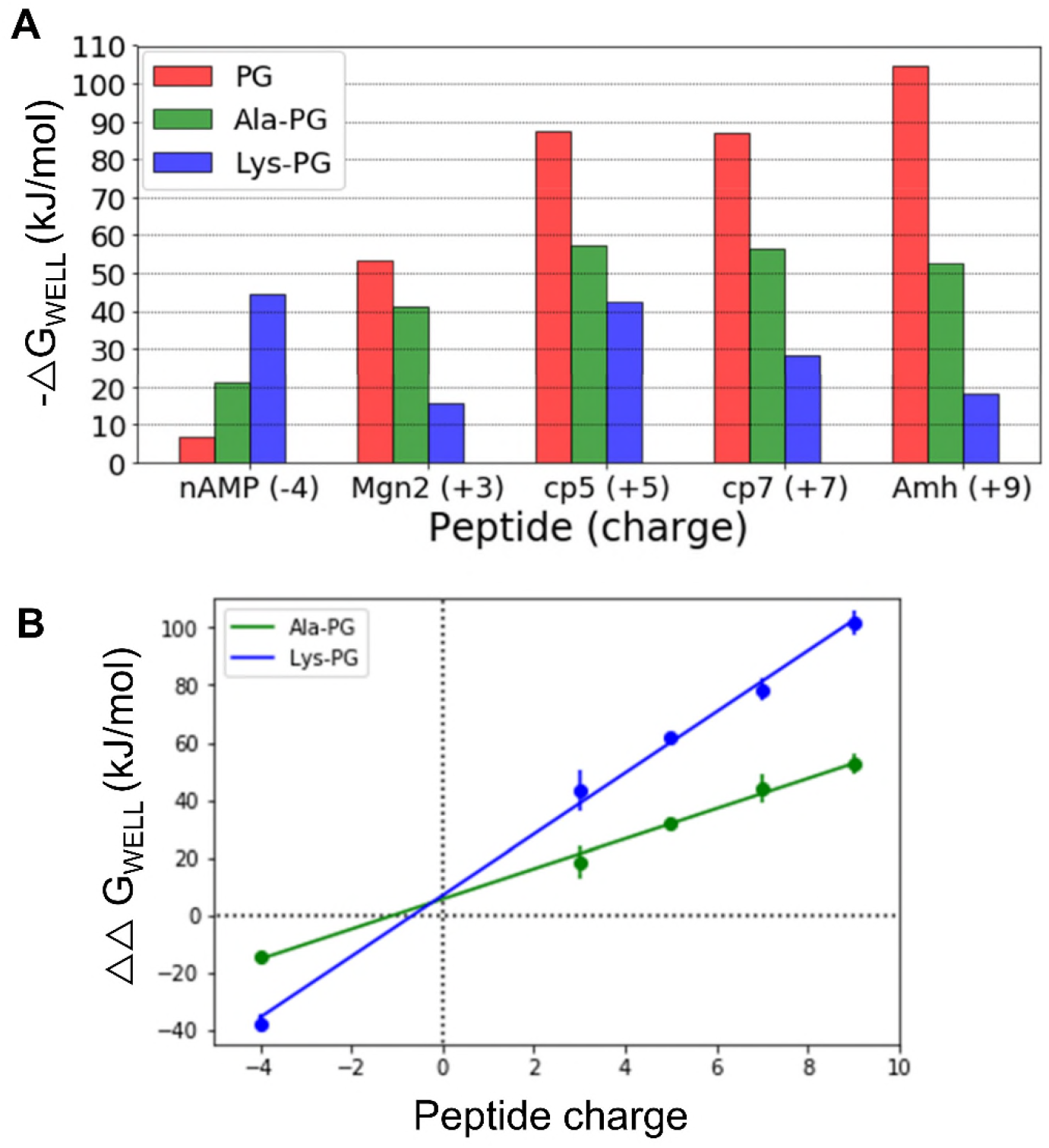
Depths of the free energy wells for peptide binding to the membrane surface derived from the PMFs shown in Fig. 5. **A** Depth of interfacial free energy wells for each peptide/membrane combination derived from the PMFs in Fig. 5. **B** Change in free energy well depth when a PG membrane is changed to either Lys-PG or Ala-PG. Graphs of data derived from **A**, showing *ΔΔG_WELL_*(PG-KPG *or* PG-APG) as a function of peptide charge with corresponding best linear fits. If *ΔΔG_WELL_* is positive that correspond to the modification of the lipid resulting in stronger binding of the peptide compared to a pure PG membrane.

All peptides show a large free energy barrier corresponding to the shift from an interfacial location to a membrane-spanning orientation. As suggested elsewhere^76^, this indicates that spontaneous insertion of single peptide molecules is unlikely to occur. However, cooperative insertion of multiple peptide molecules is likely to favour insertion and subsequent pore formation and/or membrane disruption^77^ Such peptide insertion may also be sensitive to lipid acyl tail species^78^.

## CONCLUSIONS

Our simulations clearly demonstrate that the conversion of PG into either Ala-PG or Lys-PG leads to electrostatic repulsion of positively charged peptides and a reduction in the binding of these peptides to the membrane (Figure 7). This effect is more dramatic for membrane systems containing Lys-PG, as this lipid has a greater effect on the change in the surface electrostatics of the membrane. Increasing the charge of the peptide compounds the reduction in binding. In contrast, the negatively charged peptide nAMP shows a greater binding preference for modified lipid membranes. Thus it is expected that the activity of MprF would correlate with an increase in binding of anionic peptides. However, nAMP is not a pore-forming peptide and has no antimicrobial action. However, it may provide a template for the design of peptides that can bind to and disrupt MprF-modified bacterial membranes.

**Figure 7:**
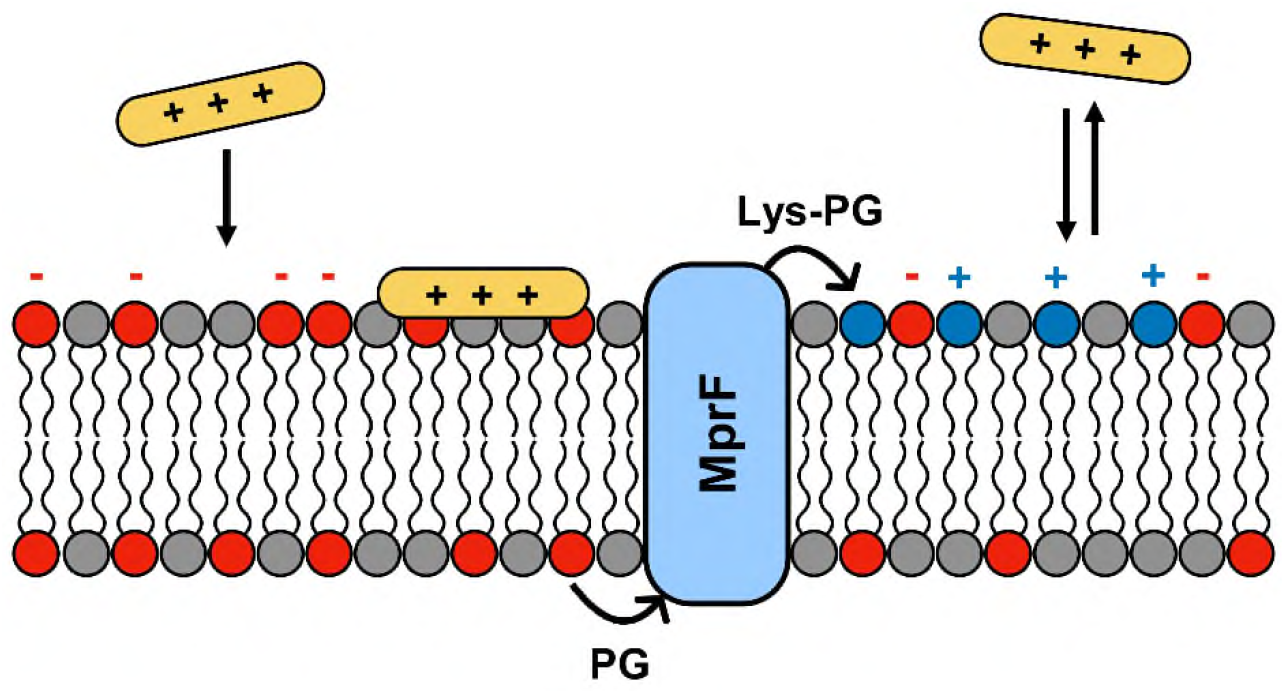
Cationic peptides (orange) bind to a negatively charged membrane. The action of the MprF protein (blue) modifies the electrostatics of the membrane via the conversion of PG lipids into Ala-PG (zwitterionic) or Lys-PG (cationic). The altered surface charge prevents peptides from binding to the membrane.

Our results from varying the bilayer compositions suggest a substantial fraction of the membrane has to be composed of Lys-PG to have a notable effect on peptide binding. From our lipid composition diagrams (Fig 4C) we see that compared to a ‘wild type-like’ membrane composed of 50:50 PG:PE, ~ 40% conversion (corresponding to reduction in membrane charge density below 0.5 *e*·nm^−2^; Fig 4D), is required to observe the beginning of a substantive drop-off in binding of cp5, cp7 and Amh. Lipid composition is variable between different species and strains of bacteria and also dependent on environmental conditions, for example Lys-PG content has been shown to increase during the growth phase^79^. Furthermore, *S. aureus* membranes can contain ~35% Lys-PG during the growth phase^32^, at which level we would predict there should be a reduction in AMP binding. At this lipid composition the MIC of the human AMP, LL-37^80^ is >300 μg/ml, compared with a *S. aureus* mutant with 0% Lys-PG in its membranes which has a MIC of ~75 μg/ml.

Undoubtedly, the mechanism of action of AMPs is much more complicated than simply binding of peptides to a bilayer. However, it is clear that if lipid modification reduces the affinity of a peptide for a bacterial membrane, then the downstream efficacy of that peptide will also be substantially reduced. It is also important to note that we investigated peptide binding to the membrane, not folding of the peptide at or close to the membrane surface. It is known that AMPs adopt their preferred secondary structure upon association with a membrane or membrane-like surface^81^. The folding process may affect how peptides penetrate the membrane and may provide a partial explanation as to why the free energy of peptides spanning the hydrophobic core is so large. This question may be addressed further using all-atomistic simulations^50^.

We can use our simulations to address the functionally relevant question of what percentage of lipid modification has to occur for AMP binding to be substantially decreased. From considerations of the computational biophysics of peptide/bilayer interactions it would seem that >40% modification may be required. As we continue to explore and design peptides with different modes of action^82-84^ in order to address antimicrobial resistance, it will become increasingly important to take account of the quantitative effects of changes in lipid composition on interactions with these peptides^85^.

## ACKNOWLEDGEMENTS

Work was funded by EPSRC in collaboration with IBM and STFC Hartree Centre’s Innovation Return on Research programme, funded by the Department for Business, Energy & Industrial Strategy. PJS and MSPS are funded by Wellcome (208361/Z/17/Z). Research in MSPS’s group is also supported by BBSRC and EPSRC. PWS is funded by EPSRC and IBM. Research in PJS’s lab is funded by the MRC (MR/S009213/1) and BBSRC (BB/P01948X/1, BB/R002517/1 BB/S003339/1). Simulations were carried out in part on ARCHER and JADE UK National Supercomputing Services, provided by HECBioSim, the UK High End Computing Consortium for Biomolecular Simulation (hecbiosim.ac.uk), which is supported by the EPSRC (EP/L000253/1).

## Notes

### Competing Interest Statement

The authors have declared no competing interest.

