## supplemental figures and tables for "Modified Bacterial Lipids Which Alter Membrane Surface Charge Reduce Binding of Antimicrobial Peptides"

### Supplementary Figures

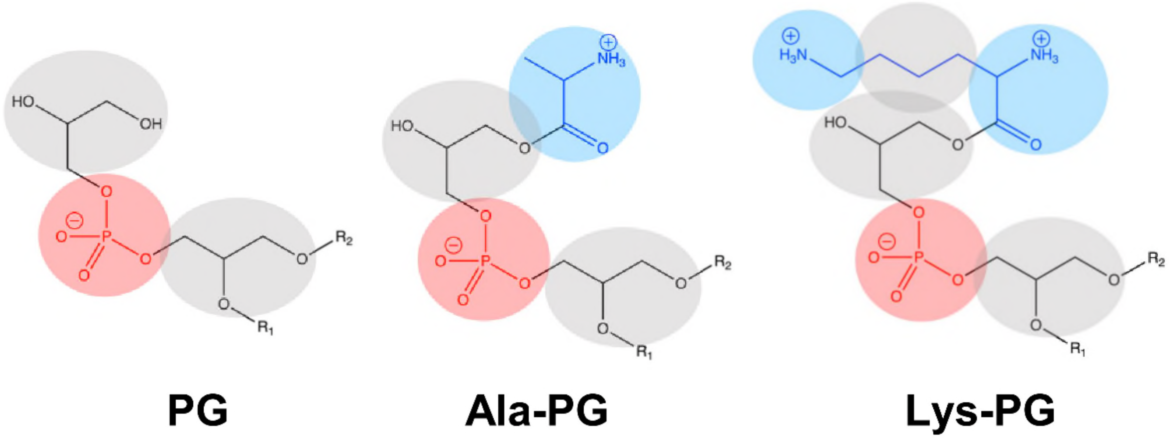


*Figure S1*

PG, Ala-PG and Lys-PG headgroups. In the Martini CG representation (coloured circles and ellipses representing Martini beads) the transformation of PG into Ala-PG corresponds to addition of a single positively charged (blue) bead. Lys-PG has three additional beads compared to PG, two of which are positively charged.

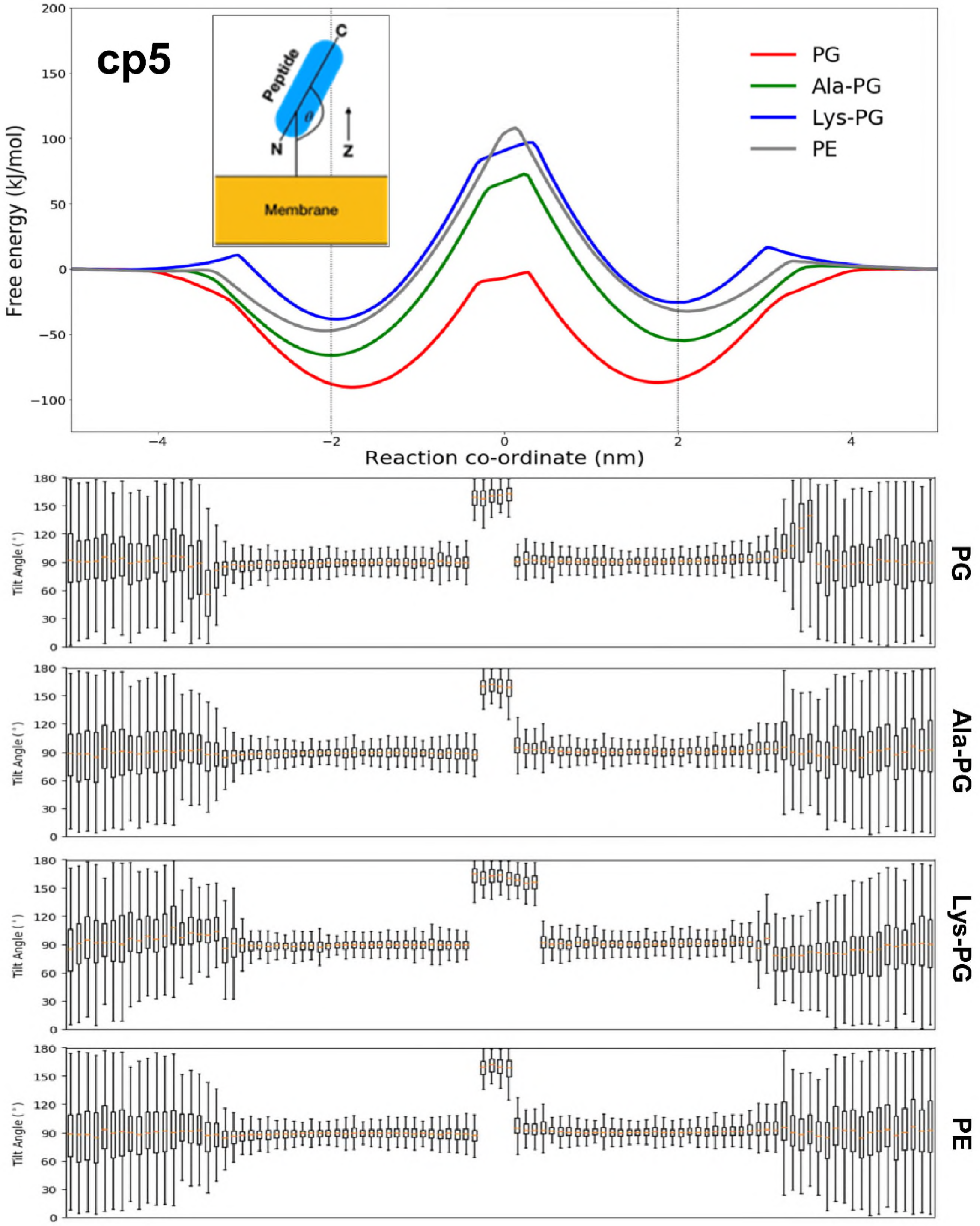


*Figure S2*

PMFs for interaction of peptide cp5 with PG, Ala-PG, Lys-PG and PE membranes (see main figure 5). The inset shows the definition of the helix tilt angle. The corresponding values of the tilt angles vs. reaction coordinate are shown below for the four different bilayers.

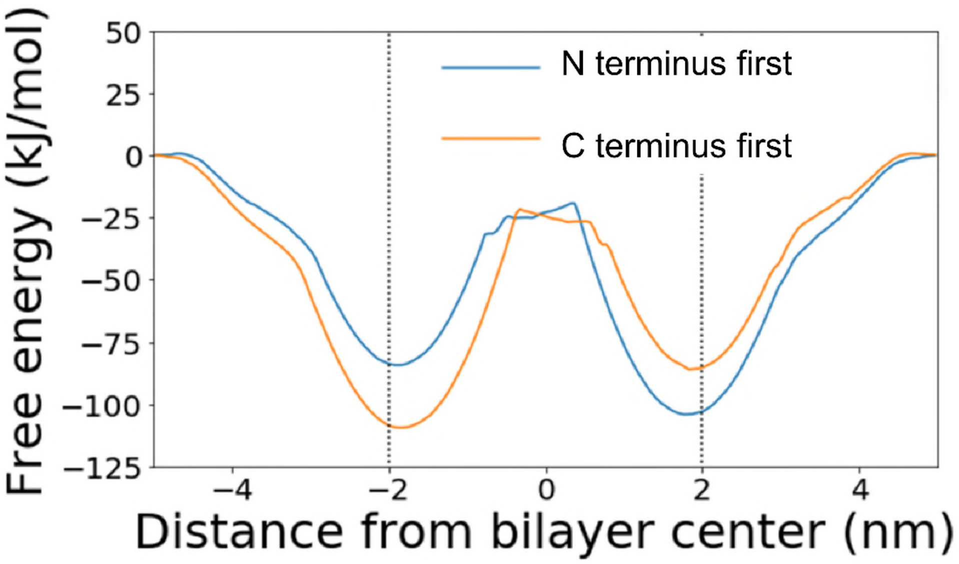


<u>*Figure S3*</u>

N terminus and C terminus first PMF landscapes for the Amh peptide and a PG membrane. PMFs for interaction of peptide Amh with a PG bilayer. In the initial steered simulation used to define the reaction coordinate either the N- (blue) or C terminus (orange) of the peptide approaches the bilayer surface first. As expected, the two resultant PMFs are seen to be approximate mirror images of one another.

**Supplementary Tables**

- Summary of Simulations Performed

*Table S1: Asymmetric bilayer systems*

| |
|---|
| upper leaflet: PG:PE (30:70) + lower leaflet: Ala-PG:PE (30:70) |
| upper leaflet: PG:PE (30:70) + lower leaflet: Lys-PG:PE (30:70) |
| 50 repeat simulations, each of duration 1 μs, were performed for each of the five peptide species. |

*Table S2: Symmetric bilayer systems*

| PE (%) | PG (%) | Ala-PG (%) | Lys-PG (%) |
|---|---|---|---|
| 100 | 0 | 0 | 0 |
| 90 | 10 | 0 | 0 |
| 90 | 0 | 10 | 0 |
| 90 | 0 | 0 | 10 |
| 80 | 20 | 0 | 0 |
| 80 | 10 | 10 | 0 |
| 80 | 10 | 0 | 10 |
| 80 | 0 | 20 | 0 |
| 80 | 0 | 0 | 20 |
| 70 | 30 | 0 | 0 |
| 70 | 20 | 10 | 0 |
| 70 | 20 | 0 | 10 |
| 70 | 10 | 20 | 0 |
| 70 | 10 | 0 | 20 |
| 70 | 0 | 30 | 0 |
| 70 | 0 | 0 | 30 |
| 60 | 40 | 0 | 0 |
| 60 | 30 | 10 | 0 |
| 60 | 30 | 0 | 10 |
| 60 | 20 | 20 | 0 |
| 60 | 20 | 0 | 20 |
| 60 | 10 | 30 | 0 |
| 60 | 10 | 0 | 30 |
| 60 | 0 | 40 | 0 |
| 60 | 0 | 0 | 40 |
| 50 | 50 | 0 | 0 |
| 50 | 40 | 10 | 0 |
| 50 | 40 | 0 | 10 |
| 50 | 30 | 20 | 0 |
| 50 | 30 | 0 | 20 |
| 50 | 20 | 30 | 0 |
| 50 | 20 | 0 | 30 |
| 50 | 10 | 40 | 0 |
| 50 | 10 | 0 | 40 |
| 50 | 0 | 50 | 0 |
| 50 | 0 | 0 | 50 |

For each of the five peptides simulations were performed with membranes of the compositions listed above, with 25 repeats (duration 1 μs) for each peptide/bilayer combination.

*Table S3: PMFs*

The following simulations were performed for each of the five peptides to generate the PMFs discussed in the main text. See the main text Methods for further details.

| Lipid bilayer | Simulation windows | Simulation duration |
|---|---|---|
| PE | 101 windows | 1 μs per window |
| PG | 101 windows | 1 μs per window |
| Ala-PG | 101 windows | 1 μs per window |
| Lys-PG | 101 windows | 1 μs per window |
